# SQUARNA: stem maximization for accurate de novo RNA secondary structure prediction

**DOI:** 10.64898/2026.06.30.735492

**Authors:** Maksim D. Serdakov, Davyd R. Bohdan, Grigory I. Nikolaev, Janusz M. Bujnicki, Eugene F. Baulin

## Abstract

Non-coding RNAs play diverse roles in a wide range of cellular processes, with their spatial structure being pivotal to their function. RNA secondary structure is a key determinant of its overall fold. Given the scarcity of experimentally determined RNA 3D structures, understanding secondary structure is vital for discerning RNA function. Currently, there is no universally effective solution for de novo RNA secondary structure prediction. Existing methods are becoming increasingly complex without marked improvements in accuracy and often overlook critical features such as pseudoknots and alternative folds. Here, we introduce SQUARNA, a new approach to de novo RNA secondary structure prediction that is suitable for both individual RNA analysis and large-scale structural searches. SQUARNA revisits the concept of base pair maximization and develops it into a stem maximization idea coupled with the widely used free energy minimization (MFE) framework. SQUARNA can predict alternative structures and handle pseudoknots of arbitrary complexity. Benchmarking shows that SQUARNA outperforms existing methods, including deep learning models, in both single-sequence and alignment-based RNA secondary structure prediction. SQUARNA seamlessly integrates sequence and alignment information with experimental data, such as residue reactivities obtained by chemical probing, as well as other structural restraints, including automated searches for Rfam database templates, G-quadruplex patterns, and protein-binding motifs. SQUARNA is available as a standalone tool at https://github.com/febos/SQUARNA and as a web server at https://larnal.imol.institute.

## INTRODUCTION

Hundreds of structured RNA molecules have been identified and characterized to date [1], ranging from ligand-binding riboswitches that regulate gene expression [2] to structured elements in viral RNAs that confer resistance to degradation [3]. The function of these RNAs is governed by their spatial structure [4], which in turn is determined by the ribonucleotide sequence and environmental conditions [5]. It is widely accepted that RNA secondary structure, consisting of canonical Watson-Crick (WC) G-C and A-U and wobble G-U base pairs, forms first, with other interactions developing subsequently [6]. Because RNA secondary structure is critical for defining the molecule’s global fold, knowledge of this structure is crucial for determining the RNA 3D structure and function [7]. Despite significant advances in experimental techniques for RNA structure determination, particularly in chemical probing methods [8] and cryo-electron microscopy (cryo-EM) [9], the structures of many functional RNAs remain unknown [10]. Consequently, the computational problem of RNA secondary structure prediction remains highly relevant.

The computational RNA secondary structure prediction problem involves determining which pairs of nucleotides form WC and wobble base pairs. Predictions can be made for a single RNA sequence or a multiple sequence alignment. Depending on the approach, the prediction may rely on known structural templates or be performed de novo. The most popular template search engine is implemented in the Infernal package [11] and is incorporated into the Rfam [10] and RNAcentral databases [12]. The most widely used approach for de novo single-sequence prediction is free energy minimization [13], in which the free energy of a structure is calculated as the sum of the free energies of its elements, and the structure with the minimum free energy (MFE) is identified using dynamic programming [14]. Some methods use probabilistic approaches to predict the structure with maximum expected accuracy (MEA) [15]. For alignment-based predictions, a classical approach involves identifying covarying residue pairs that correspond to evolutionary conserved base pairs [16]. Several methods combine covariation analysis with algorithms to find either the MFE [17] or the MEA structure [15, 16, 18]. More recently, deep learning methods have been developed for both single-sequence and alignment-based prediction, demonstrating superior performance compared to traditional tools [19–22].

Despite the diversity of approaches, there is still no definitive solution for RNA secondary structure prediction, particularly for single RNA sequences or sequence alignments containing too few or too similar sequences for effective covariation analysis [23]. Methods are becoming increasingly complex without notable improvements in accuracy [21]. Most methods ignore pseudoknots [18, 19, 24], which represents a significant oversimplification [25]. In addition, the majority of methods predict only a single structure and are therefore unsuitable for RNA molecules that adopt alternative conformations [15, 19, 20, 24], with only a few exceptions [24, 26]. Many tools cannot incorporate structural restraints or chemical probing data into their predictions [15, 16, 18–20]. Only a limited number of tools predict structures formed by multiple RNA sequences [27]. With rare exceptions [28], existing tools consider only “classical” secondary structures, requiring researchers to use separate tools to explore non-canonical formations such as G-quadruplexes [29]. Finally, deep learning methods often suffer from overfitting, lack interpretability, and do not generalize well to RNAs that exhibit previously unseen structures [30, 31].

In this work, we present SQUARNA, a new RNA secondary structure prediction method based on a stem maximization model that addresses the limitations of previous approaches. Benchmarking demonstrates that SQUARNA outperforms existing tools for both single-sequence and sequence alignment inputs. SQUARNA seamlessly incorporates structural restraints, including chemical probing data, and supports large-scale structural searches through a flexibly configurable implementation. The web server further enhances SQUARNA’s accessibility and enables automated searches for Rfam templates, G-quadruplex patterns, and protein-binding motifs, providing a universal entry point for RNA structure analysis.

## RESULTS

### Stem maximization concept

We formulate the de novo RNA secondary structure prediction problem for a single-sequence input as a variant of the Maximum Weighted Matching (MWM) problem, wherein a set of nodes (residues) and a set of weighted edges between them (possible base pairs) define the input graph, and an optimal matching between selected residues constitutes the output [32]. We represent the input graph as an adjacency matrix, where each cell *(i, j)* stores the edge weight corresponding to the base pair formed between residues *i* and *j*, with positive weights assigned to G-C, A-U, and G-U pairs and zero weights assigned to all others (Figure 1A). To account for base pair symmetry (if *i* interacts with *j*, then *j* interacts with *i*) and the fact that RNA does not form hairpins shorter than three residues [33], only edges satisfying *j > i + 3* are considered.

**Figure 1.**
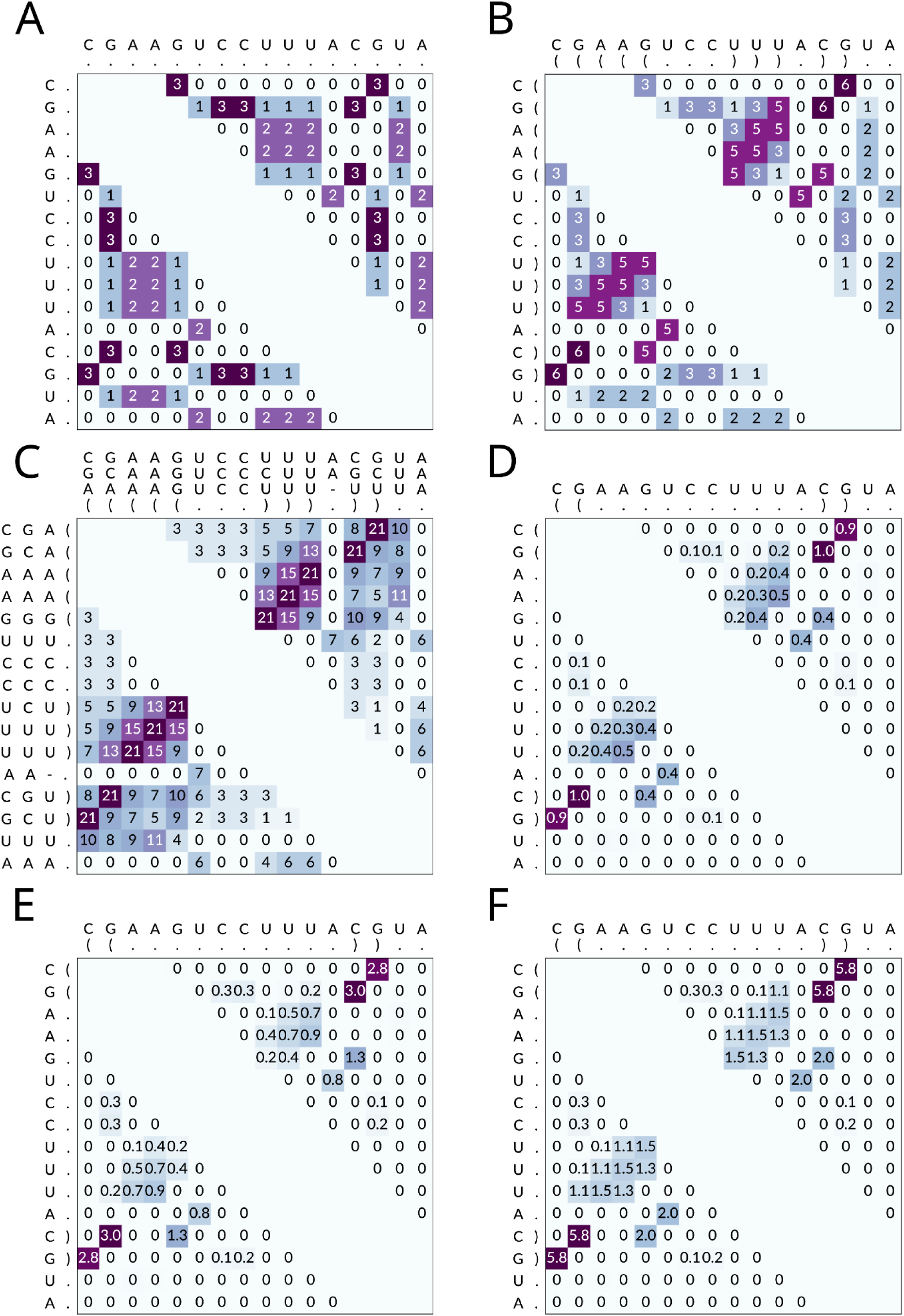
Examples illustrating base pair and stem weight matrices. (**A**) Base pair weight matrix with weights of 3, 2, and 1 assigned to G-C, A-U, and G-U base pairs, respectively. (**B**) Stem weight matrix derived from the matrix in panel A by summing the values along consecutive non-zero diagonals. (**C**) Stem weight matrix calculated for a sequence alignment as the sum of three individual stem weight matrices. Each individual matrix is computed for an ungapped sequence and subsequently realigned. (**D**) Base pair probability matrix generated by the ViennaRNA package [24]. The matrix is normalized by dividing each value by the maximum probability. (**E**) Product of the matrices shown in panels A and D. (**F**) Stem weight matrix derived from the matrix in panel E by summing the values along consecutive non-zero diagonals. In all panels, only base pairs (i, j) with j > i + 3 and their symmetric (j, i) values are shown.

Recognizing the interdependence of base pairs [34, 35] and the tendency of base pairs *(i, j)* to co-occur with neighboring base pairs *(i + 1, j - 1)* and *(i - 1, j + 1)*, we use the weights of non-extendable stems (maximal stacks of consecutive base pairs, represented as continuous non-zero diagonal stretches in an adjacency matrix) as weights for potential matchings to guide the predictions (Figure 1B, 1F). The *stem weight* is defined as the sum of the weights of its constituent base pairs. The use of only non-extendable stems, without considering all possible partial formations, is justified by the observation that nearly 87% of stems in the PDB database [36] either correspond to fully formed non-extendable stems or lack at most one flanking base pair (Supplementary Table S1). Notably, a stem weight matrix can be naturally obtained for a sequence alignment by summing the matrices of individual sequences (Figure 1C). We therefore formulate the *stem maximization concept* as a variant of base pair maximization in which base pairs are scored according to stem-based weights, and the structure that maximizes the total score is considered the optimal weighted matching. For base pair scoring, we consider a combination (either a sum or a product) of two values (Figure 1A, 1D, 1E), see the Methods section for details. The first value is a fixed base pair weight that depends on the identities of the paired nucleotides. The default weights used in SQUARNA are 3.25 for G-C, 1.25 for A-U, and -1.25 for G-U base pairs. The second value is the base pair probability derived by the ViennaRNA package [24] within the MFE framework.

To solve the formulated MWM problem, i.e., to convert stem weight matrices into RNA secondary structures, we developed SQUARNA (SQUAre RNA), which incorporates four independent algorithms: Nussinov [37], Hungarian [38], Edmonds [32, 39], and an original greedy approach, see the Methods section for details. Interestingly, unlike the original Nussinov approach, in which all base pairs are assigned a weight of 1 [37], the variant using our stem weight matrices, even without the base pair probability component, successfully predicts the canonical cloverleaf structure of tRNAs (Supplementary Figure S1). Furthermore, the Edmonds algorithm has been shown to predict RNA secondary structures with pseudoknots in polynomial time [32], whereas this problem is known to be NP-hard under the MFE framework [40, 41]. Thus, the proposed stem maximization concept and its implementation in SQUARNA address both single-sequence and alignment-based RNA secondary structure prediction, enabling the prediction of alternative conformations and the handling of pseudoknotted structures through the use of multiple complementary algorithms.

### Single-sequence prediction with SQUARNA

To identify optimal parameters for SQUARNA and benchmark its performance, we prepared two manually curated datasets of RNA sequences derived from PDB entries: a training dataset (SRtrain, 274 sequences) and a test dataset (SRtest, 247 sequences), with sequence similarity below 80% between the two datasets, see the Methods section for more details. The F-score was selected as the target metric, and prediction quality was assessed using either the top-ranked prediction or, where applicable, the best of the top five predictions generated by each method. Considering the top five predictions is justified by the fact that RNA 3D structure prediction competitions, such as RNA-Puzzles [42] and CASP [43], allow up to five predictions per target. The optimized version of SQUARNA was effective in identifying alternative conformations. For example, among its top five predictions for the S15 leader sequence [44] (Figure 2C), the two known conformations that regulate expression of the S15 ribosomal protein were ranked first (consecutive hairpins) and third (pseudoknot).

**Figure 2.**
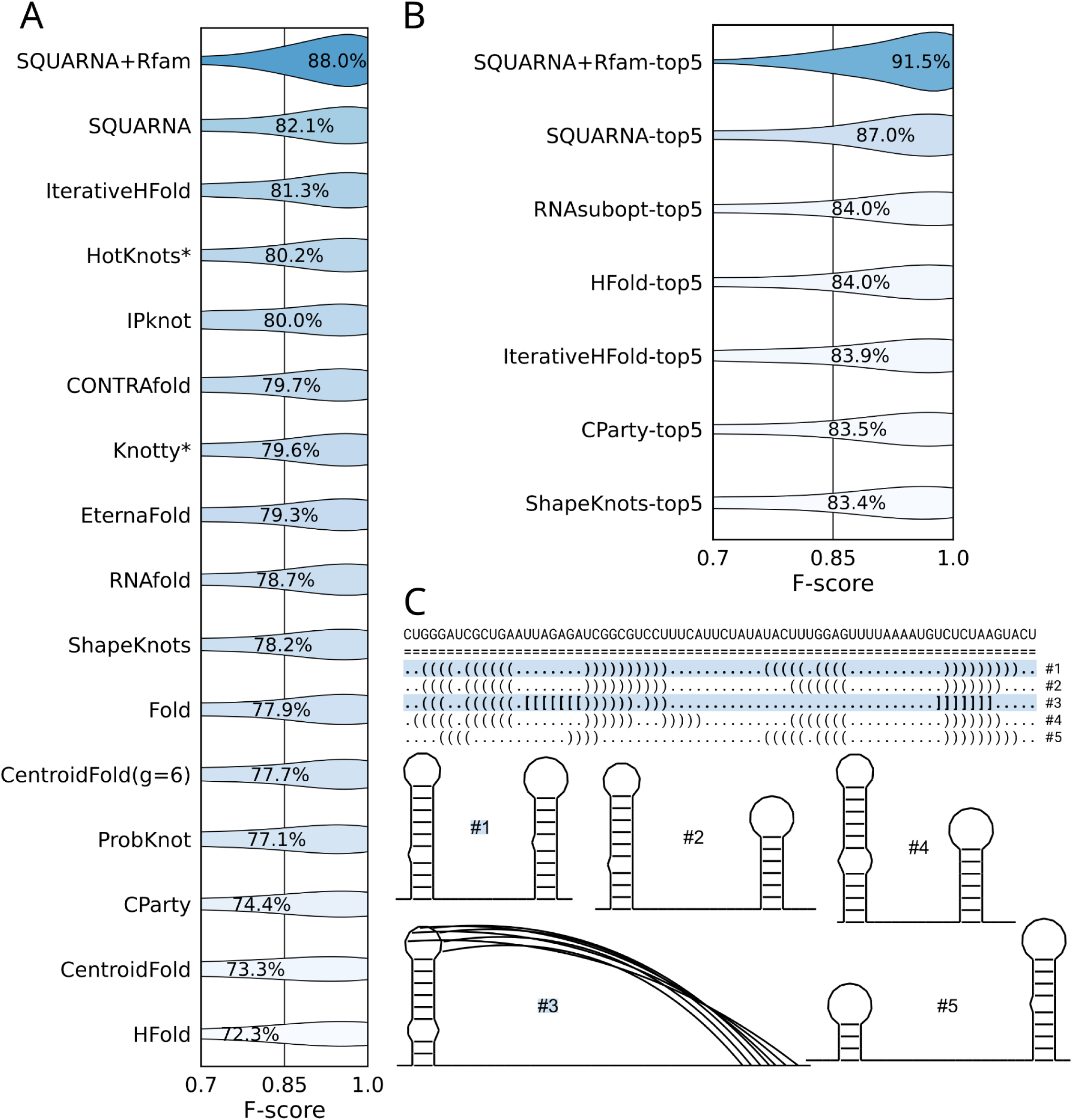
Single-sequence SQUARNA performance. (**A**) Violin plot of F-score values achieved by single-sequence predictors on the SRtest dataset (247 sequences), considering only the top-ranked prediction. (**B**) Violin plot of F-score values achieved by single-sequence predictors on the SRtest dataset (247 sequences), considering the best of the top five predictions. Mean value is labeled on each violin. Asterisks denote tools that could not process all sequences in the dataset and were assigned an F-score of 0.0 for those sequences, see the Methods section for details. (**C**) Top five secondary structures predicted with SQUARNA for the S15 leader sequence [44]. The experimentally confirmed conformations, ranked #1 and #3, are highlighted with a blue background and bold font. Structure diagrams were prepared using RNArtist (https://github.com/fjossinet/RNArtist).

We compared SQUARNA with existing single-sequence predictors [15, 19, 20, 22, 24, 45–55] on the SRtest dataset. SQUARNA achieved the highest mean F-score of 82.1% for top-one predictions (Figure 2A) and the highest mean F-score of 87.0% when considering the best of top five predictions (Figure 2B). In addition, we implemented a hybrid mode (SQUARNA+Rfam), in which Rfam template search powered by cmscan [11] precedes prediction. In this mode, the identified templates serve as structural restraints for subsequent prediction. SQUARNA+Rfam reached 88.0% F-score for top-one predictions (Figure 2A) and 91.5% F-score for top-five predictions (Figure 2B). For additional reference, we also benchmarked deep learning models (Supplementary Figure S2A) and observed that only one recent model, RibonanzaNet-SS, matched the performance of SQUARNA+Rfam with an F-score of 87.9%. However, RibonanzaNet-SS was fine-tuned on PDB structures [54], and thus data leakage is expected. Overall, SQUARNA demonstrates state-of-the-art performance while avoiding the data-related limitations of artificial intelligence-based approaches.

We analyzed the contributions of the SQUARNA components (Supplementary Figure S2B, S2C). The Hungarian and Edmonds algorithms were each sufficient on their own to reach the best SQUARNA performance in top-one predictions, whereas the greedy algorithm was the key contributor for top-five predictions. Base pair probabilities derived from ViennaRNA on average contributed 4-7 percentage points to SQUARNA performance. Additionally, alongside Rfam template search, we implemented options for G-quadruplex pattern recognition and protein-binding motif annotation, see the Methods section for details. All three options are enabled by default in the web server implementation (https://larnal.imol.institute/), which is designed specifically for the analysis of individual RNA sequences.

### Alignment-based prediction with SQUARNA

For alignment-based RNA secondary structure prediction, the greedy algorithm with base pair scores of 3.25 for G-C, 2.00 for A-U, and -1.00 for G-U, and without a base pair probability component, demonstrated the best performance, see the Methods section for details. These parameters were defined using the RNAStralignExt dataset (Supplementary Table S2), which we compiled from eight Rfam families [10] in the RNAStralign dataset [56], together with three additional families with well-characterized pseudoknotted structures: HDV ribozyme, SAM riboswitch, and Twister ribozyme, included to improve pseudoknot representation.

We benchmarked SQUARNA for alignment-based prediction against existing methods [15–18] on the full set of Rfam 14.9 seed alignments, comprising 4,108 non-coding RNA families (Figure 3A). For alignments containing 50-300 sequences, all predictors showed comparable performance, whereas SQUARNA and R-scape performed better on deeper alignments, and CentroidAlifold and RNAalifold performed better on shallow alignments of 2-30 sequences. The latter can be explained by the fact that shallow Rfam alignments are often annotated with predicted structures; for example, 66% of seed alignments of 2-4 sequences feature secondary structures predicted by RNAalifold (Supplementary Table S3). On the RfamPDB dataset, which includes 134 families represented with 3D structures in the PDB database [36], SQUARNA matched the performance of MFE-based tools on shallow alignments and showed the best performance for alignments of more than 200 sequences (Figure 3B).

**Figure 3.**
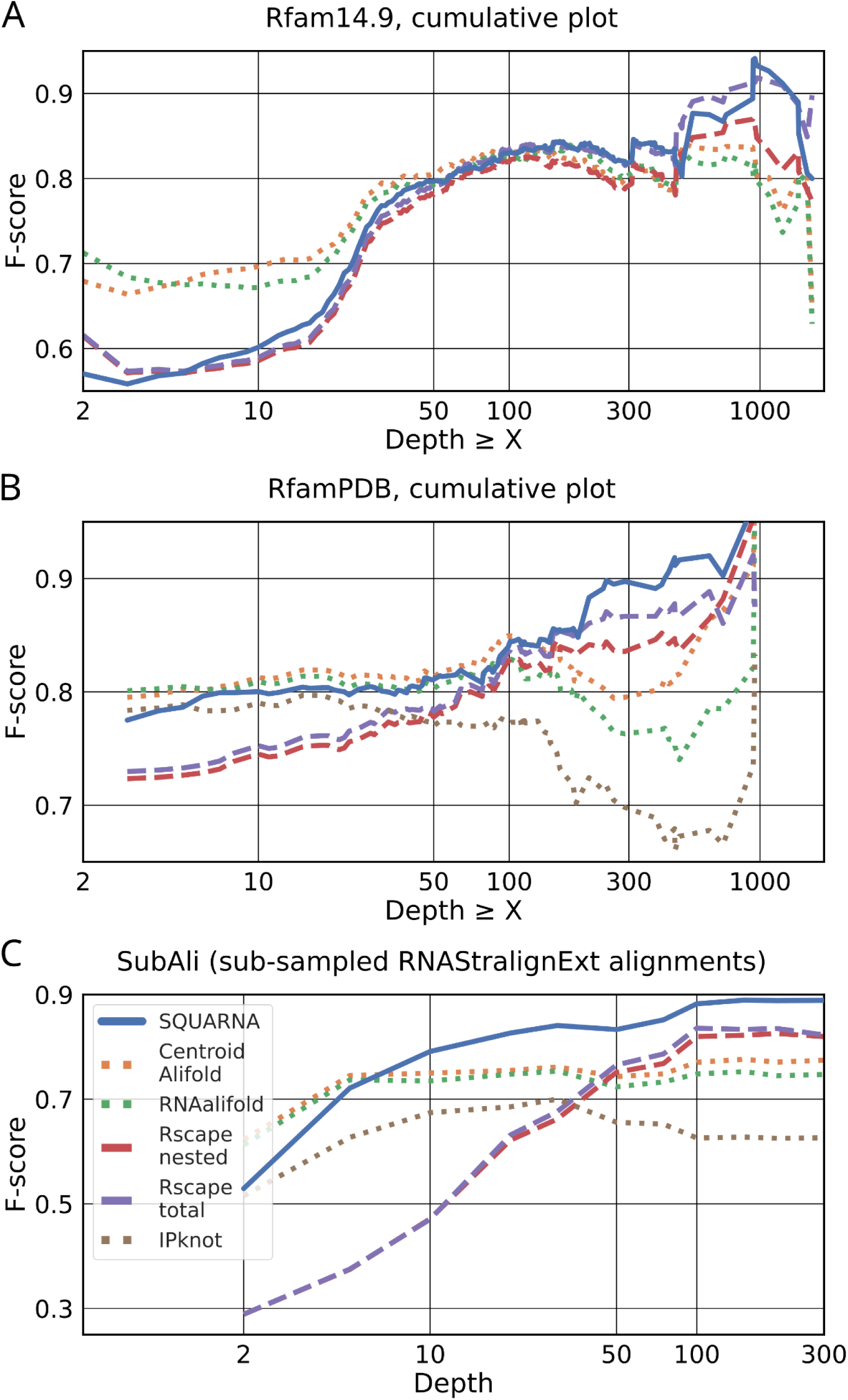
Alignment-input SQUARNA performance. (**A**) Cumulative plots of prediction F-scores against alignment depth threshold, as demonstrated by alignment-based predictors on the Rfam 14.9 dataset of 4,108 multiple sequence alignments. We failed to obtain IPknot predictions for all families of the dataset and omitted it from the benchmark. (**B**) Cumulative plots of prediction F-scores against alignment depth threshold, as demonstrated by alignment-based predictors on the RfamPDB dataset of 134 multiple sequence alignments with experimentally confirmed secondary structures. (**C**) Performance of alignment-based predictors on the SubAli dataset of sub-sampled alignments from the RNAStralignExt families.

To mitigate bias arising from the uneven distribution of alignment depths across Rfam families, we compiled the SubAli dataset of randomly sub-sampled alignments with depths ranging from 2 to 300 sequences for each family in RNAStralignExt. On this dataset, SQUARNA outperformed all other tools (Figure 3C), and only SQUARNA and R-scape showed consistent performance improvements with increasing alignment depth (Supplementary Figure S3). SQUARNA started with a 50% mean F-score for alignments of two sequences, reached 80% at ten sequences, and stabilized at approximately 90% for alignments exceeding 100 sequences. CentroidAlifold and RNAalifold started with a mean F-score of 60% for two sequences and plateaued at around 75% from five sequences onward. R-scape exhibited a sharp increase from 30% to 80-85% mean F-score between 2 and 100 sequences, stabilizing thereafter. To assess the influence of sequence similarity on prediction quality, we constructed the SeqSim dataset by randomly sampling 5-, 10-, and 20-sequence alignments with varying mean pairwise sequence similarity across families from RNAStralignExt. Interestingly, only SQUARNA exhibited a subtle yet discernible negative correlation between prediction quality and mean pairwise sequence similarity in the alignment, which diminished with increasing alignment depth (Supplementary Figure S4). This negative correlation underscores the method’s ability to utilize the signal from more diverged (less similar) sequences.

We then examined specific instances in which SQUARNA encountered difficulties across the Rfam 14.9 dataset. For the ribosomal S15 leader sequence (RF00114), SQUARNA achieved an F-score of only 32%, as it predicted the correct pseudoknot conformation, whereas the Rfam-annotated structure corresponds to the alternative fold of two consecutive hairpins [44]. As this example illustrates, secondary structures currently annotated in Rfam are not always an ideal ground truth. Therefore, as an additional test, we assessed the number of base pairs predicted by SQUARNA in the Rfam 14.9 seed alignments that were also identified as significantly covarying by R-scape [57]. SQUARNA predictions included 5,540 covarying base pairs, comparable to the 5,535 base pairs in the Rfam-annotated secondary structures, whereas CentroidAlifold and RNAalifold each identified approximately 5,000 covarying base pairs (5,016 and 4,989, respectively; Supplementary Figure S5). Thus, SQUARNA-derived secondary structures are supported by R-scape-identified covariation to a degree comparable to Rfam annotations and higher than that of other methods.

### Chemical probing input data

SQUARNA was implemented to integrate user-specified chemical probing data in the form of individual residue reactivities. The input reactivities were used independently in the calculation of two base pair score components: the base-type-specific weights (bpw, custom formula) and the ViennaRNA-derived base pair probabilities (bpp, default ViennaRNA method for SHAPE data [58]), see the Methods section for details. To assess the impact of chemical probing data on SQUARNA prediction quality, we used the S01 dataset of 24 challenging RNAs with SHAPE data [26]. Using only sequence input, SQUARNA achieved a mean F-score of 68.7% for top-ranked predictions and 76.6% for the best of the top five predictions (Figure 4). The default SQUARNA configuration with chemical probing data demonstrated substantially improved performance, reaching 77.9% and 84.6% F-score for top one and top five predictions, respectively. Notably, incorporating residue reactivities into each of the two base pair score components individually improved prediction quality to a comparable extent, with the bpp component dominating top one predictions, whereas the bpw component had a greater impact on top five predictions (Figure 4).

**Figure 4.**
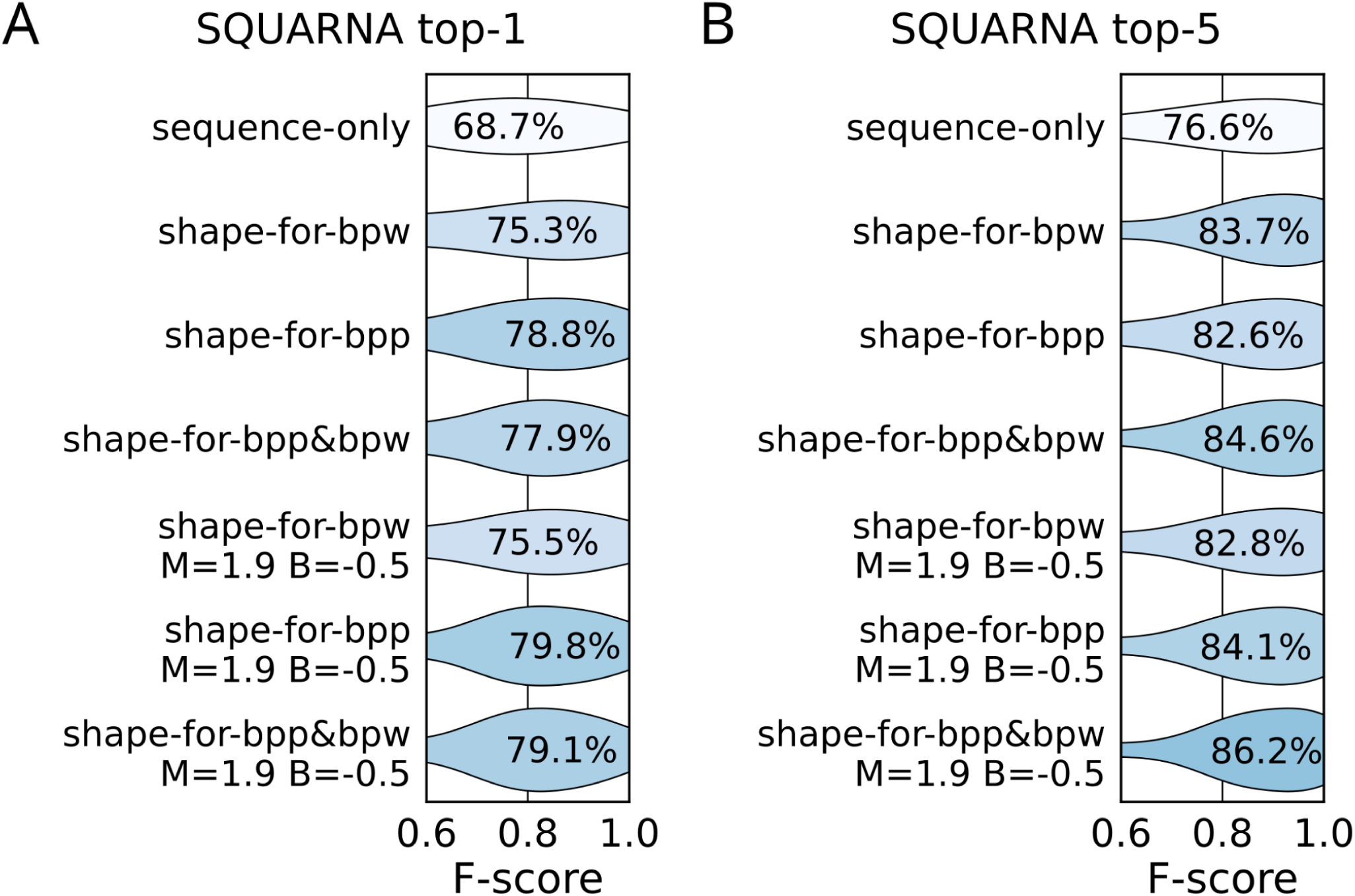
SQUARNA performance with input residue reactivities. F-score distributions for (**A**) top-ranked predictions and (**B**) the best of the top five predictions generated by SQUARNA for the 24 RNA sequences in the S01 dataset. Mean value is labeled on each violin. The “sequence-only” variant does not use residue reactivities and relies solely on sequence input; “shape-for-bpw” uses reactivities only for the base pair weight component of the SQUARNA scoring matrices; “shape-for-bpp” employs ViennaRNA functionality to incorporate reactivities, affecting only the base pair probability component of the scoring matrices; “shape-for-bpp&bpw” indicates the default SQUARNA configuration; the *“M=1.9 B=-0.5”* variants show SQUARNA performance using optimized M and B values obtained for the S01 dataset through grid search.

We also identified optimal slope (M = 1.9) and intercept (B = -0.5) parameters [58] through grid search on the S01 dataset, resulting in a further improvement of approximately 1-1.5 percentage points. For additional reference, we benchmarked SQUARNA against existing methods capable of incorporating chemical probing data, namely RNAfold/RNAsubopt [24] and ShapeKnots [26]. SQUARNA performed on par with RNAfold for top-one predictions and slightly outperformed RNAsubopt when considering the best of the top five predictions, whereas ShapeKnots, which was originally trained on a subset of the S01 dataset, achieved the highest performance in both settings (Supplementary Figure S6). Overall, SQUARNA effectively utilized input residue reactivities, achieving prediction quality comparable to that of state-of-the-art methods.

## DISCUSSION

In this work, we introduced SQUARNA, a new approach to de novo RNA secondary structure prediction based on the stem maximization framework. Our benchmarks demonstrate that SQUARNA outperforms existing methods, including deep learning models, in both single-sequence and alignment-based prediction tasks. Furthermore, SQUARNA was shown to efficiently incorporate additional input data, such as residue reactivities derived from chemical probing experiments. SQUARNA was implemented both as a standalone tool suitable for large-scale structural searches and as a web server for the analysis of individual RNAs, including automated annotation of Rfam templates, G-quadruplexes, and protein-binding motifs.

The state-of-the-art performance of SQUARNA demonstrates that non-extendable stems provide a rich source of additional signal that can be leveraged in base pair scoring. To the best of our knowledge, despite the high abundance of such stems in experimentally determined RNA 3D structures (Supplementary Table S1), their use has been largely limited to kinetics-based approaches, such as the “saturated helices” employed in a cotranscriptional folding model [59]. However, a concept similar to the stem score matrix, termed a “helix plot” and described as “sort of a stacking energy derived from helices” was introduced by Tabaska et. al [32] in 1998. In that work, the authors presented a polynomial-time method for RNA secondary structure prediction with pseudoknots based on the Gabow implementation [60] of the Edmonds algorithm for the MWM problem [39]. Unfortunately, mutual information values were used as base pair scores, limiting the applicability of the method to sequence alignments. At the same time, MFE-based methods were later shown to be NP-hard in the presence of pseudoknots [40, 41], and their success contributed to the widespread misconception that pseudoknot prediction itself is NP-hard [61, 62]. Overall, we believe that the MWM framework, together with its polynomial-time algorithms, remains underappreciated in the field of RNA structure prediction, and that the development of improved base pair scoring schemes represents a promising direction for future research.

While single-sequence SQUARNA outperformed existing conventional tools, several deep learning models still achieved better prediction accuracy. Only when coupled with a preceding Rfam template search using cmscan [11] did SQUARNA surpass AI-based methods. An important aspect of such benchmarks is the question of how fair it is to compare conventional de novo predictors with AI approaches. Although such comparisons are reasonable from the end-user perspective, it is nearly impossible to construct a test set that completely eliminates the effects of data leakage. In this context, the SQUARNA+Rfam prediction mode may serve as a useful baseline for future benchmarking of AI-based methods. Additionally, because SQUARNA is not a deep learning model, its parameter optimization and benchmarking could be performed using high-quality datasets derived from RNA 3D structures deposited in the PDB, avoiding the need for large datasets such as bpRNA-1m [63], which include, for example, RNAalifold-predicted structures derived from Rfam [10] (Supplementary Table S3). Likewise, the 80% sequence similarity threshold between the SRtrain and SRtest datasets, while appropriate for tuning the relatively small number of SQUARNA parameters, would be insufficient to prevent data leakage during the training of AI-based models. Thus, SQUARNA achieved the performance level of leading deep learning methods while employing a substantially simpler approach that avoids common limitations of AI algorithms, including overfitting, lack of interpretability, and poor generalization to previously unseen data [30]. Moreover, the state-of-the-art performance of SQUARNA has been confirmed by independent benchmarks [64].

Alignment-based SQUARNA benefited most from the adoption of stem scores, which enhanced the signal-to-noise ratio and enabled it to reach the CentroidAlifold/RNAalifold baseline with as few as five sequences, whereas R-scape required approximately 50 sequences to achieve comparable performance (Figure 3C). Notably, a substantial fraction (46.5%) of Rfam seed alignments contain 5-50 sequences, whereas only 7.7% contain more than 50 sequences, a depth sufficient for R-scape to confidently predict a structure. Overall, SQUARNA represents a versatile solution for RNA secondary structure prediction, applicable to a broad range of practical tasks, including single-sequence prediction, alignment-based prediction, prediction with hard and soft structural restraints such as forced base pairs and residue reactivities, prediction of secondary structures shared by multiple RNA sequences, and automated identification of structural restraints such as Rfam templates, G-quadruplexes, and protein-binding motifs. Furthermore, SQUARNA handles pseudoknots of arbitrary complexity and can report alternative conformations, although, to the best of our knowledge, no suitable benchmark dataset is currently available to systematically evaluate the latter capability.

The major limitations of the current SQUARNA implementation include its relatively slow performance compared with tools from the ViennaRNA package, its reliance on ViennaRNA-derived base pair probabilities, which are biased against pseudoknots, and its dependence on alignment quality for alignment-based predictions. The first two limitations will be addressed in future versions of SQUARNA, whereas the third is planned to be overcome through an independent tool that is currently under development. In addition, we plan to extend SQUARNA to support non-canonical base pairs and modified residues, while continuing to develop improved base pair scoring schemes. The web server implementation will also be extended with complementary solutions, including tools for RNA 3D structure prediction that will incorporate SQUARNA as part of the analysis pipeline. The high performance of SQUARNA, together with its versatile implementation and broad accessibility, will greatly facilitate large-scale RNA structural screens, which form the foundation of modern efforts to discover new drugs and develop RNA-based therapeutics.

## MATERIALS AND METHODS

### Theoretical background

The SQUARNA approach is rooted in the ARTEM algorithm, which is designed for the superposition of two arbitrary RNA 3D structure fragments without prior knowledge of nucleotide matchings between the fragments [65]. The two core ideas of ARTEM are the formulation of the problem as a partial assignment problem and the utilization of dependencies in the data. In the context of de novo RNA secondary structure prediction, the partial assignment pertains to base pairs formed between nucleotides, while the primary data dependency is the tendency of base pairs to form continuous stems.

In the initial phase of our preliminary analysis, we observed optimal results with negative weights for wobble G-U base pairs. This introduced an ambiguity in the definition of a stem: either as the longest sequence of consecutive base pairs or as the subsequence of consecutive base pairs with the highest total weight. Stems in known RNA secondary structures were found to predominantly correspond to the longest stems, lacking at most one flanking base pair (Supplementary Table S1). We tested all possible stem definitions, including defining a stem as any subsequence of consecutive base pairs. However, the results remained largely unchanged. Consequently, we defined a stem as the longest possible sequence of consecutive base pairs, as this proved most efficient in terms of algorithmic runtime.

For the description of RNA secondary structure elements, such as loops and pseudoknots, we used the definitions from [66]. *Pseudoknot order* (or *pseudoknot complexity*) was defined as the minimum number of distinct bracket types required for its non-ambiguous representation in dot-bracket format. Bracket-type assignment for the base pairs of a given structure was performed greedily, with a pre-sorting of base pairs in increasing order of the number of overlaps, minimizing the number of higher-order base pairs (see the PairsToDBN function in the code: https://github.com/febos/SQUARNA/blob/main/src/SQUARNA/SQRNdbnseq.py). Two base pairs *(i, j)* and *(k, l)* were considered *inconsistent* (*intersecting*) if they could not be formed simultaneously (i.e., if *i = k* or *i = l* or *j = k* or *j = l*), and *consistent* (*non-intersecting*) otherwise. Two stems were considered *consistent* if all their base pairs were consistent; otherwise, the stems were considered *intersecting*. We defined a *base pair weight* as a fixed value depending on nucleotide types (W_GC_, W_AU_, W_GU_), a *base pair probability* as the corresponding value from the ViennaRNA-derived base pair probability matrix [24], and a *base pair score* as either the product or the sum (depending on the config, see the algorithm below) of its weight and probability. A *stem score* was defined as the sum of the scores of its base pairs.

### SQUARNA algorithm for single-sequence prediction

For the custom greedy approach, SQUARNA employs a two-step procedure. First, the pair *(i, j)* with the highest stem score is selected and marked as a base pair, and the scores are then recalculated by removing incompatible base pairs *(i, k ≠ j)* and *(m ≠ i, j)* from further consideration. This procedure is iteratively repeated until no matches above the defined threshold remain. Since all pairs within a stem have, by definition, identical stem scores, we reformulated the approach into an equivalent but more straightforward stem-based representation. In this formulation, all potential stems are enumerated and assigned scores; the stem with the highest score is selected, incompatible base pairs are removed, and the procedure is iterated. If multiple incompatible stems with comparable scores (within a defined threshold) are selected, the algorithm branches and proceeds independently for each case, with each selected stem added to the current secondary structure. Consequently, SQUARNA generates a set of alternative structures. To better capture intrinsic properties of RNA secondary structure [33], the greedy approach uses *adjusted stem scores* rather than raw stem scores, with adjustment recalculated at each iteration. The adjusted stem score is defined as the product of the stem score and three factors dependent on the previously predicted stems:

- a distance factor related to the number of residues confined by the stem, excluding residues confined by nested stems [66]:

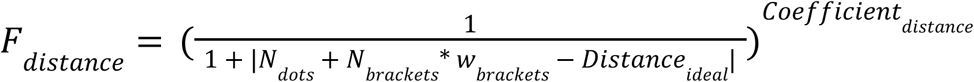

where *N_dots_*is the number of unpaired residues, *N_brackets_* is the number of paired residues, *w_brackets_* is the weight assigned to paired residues (configurable parameter *bracketweight*), *Distance_ideal_*is the “perfect” loop distance, and *Coefficient_distance_* is the exponent coefficient (configurable parameter *distcoef*).

- a pseudoknot factor related to the order of the potential pseudoknot:

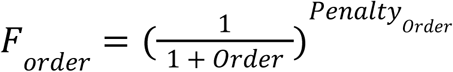

where *Order* is the minimum number of unique bracket types required to represent the given stem and the previously predicted stems in a dot-bracket diagram, and *Penalty_Order_*is the corresponding coefficient (configurable parameter *orderpenalty*).

- a loop factor providing a bonus for either a GNRA tetraloop or a short near-symmetric internal loop:

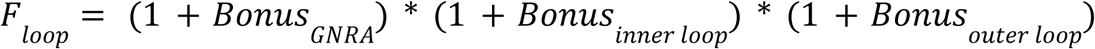

where *Bonus_GNRA_*is a bonus for GNRA tetraloop confined by the given stem, and *Bonus_inner loop_* and *Bonus_outer loop_* are equal bonuses for short near-symmetric internal loops inside and outside the given stem (configurable parameter *loopbonus*).

The single-sequence SQUARNA algorithm comprises the following steps:

0) Calculate base pair scores based on the base pair weights and base pair probabilities (configurable parameters bpweights[GC], bpweights [*AU], bpweights[GU],* and *bpp*):

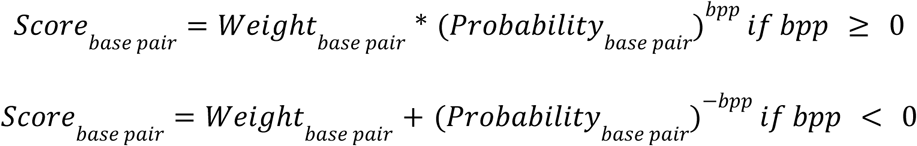

where *Probability*_*base pai*_ values are calculated using ViennaRNA [24] with [RNA.fold_compound.bpp()](https://www.tbi.univie.ac.at/RNA/ViennaRNA/doc/html/api_python.html).

Calculate the initial stem score matrix based on the base pair scores, the minimum stem length threshold (configurable parameter *minlen*) and the minimum stem score threshold (configurable parameter *minbpscore*). Predict structures using the specified global optimization algorithms (configurable parameter *algorithms*) based on the prepared stem score matrix. The available algorithms include Nussinov (N) [37], Hungarian (H) [38], and Edmonds (E) [39, 60]. If specified, proceed with the original greedy approach (G, steps 1-7), otherwise go to step 8;

1. Initialize an empty secondary structure as the only currently predicted structure;
2. Annotate all possible stems that exceed the minimum stem length and stem score thresholds and are consistent with, but not a part of, the currently predicted structure;
3. Calculate adjusted stem scores and filter out stems below the minimum adjusted stem score threshold (the product of the configurable parameters *minbpscore* and *minfinscorefactor*);
4. Sort the stems in decreasing order of their adjusted stem scores;
5. Select the highest-scoring stem along with all incompatible stems whose scores are within the suboptimality threshold (configurable parameter *subopt*);
6. Add each of the selected *k* stems to the currently predicted structure separately, resulting in *k* new currently predicted structures;
7. Repeat steps 2-6 for each currently predicted structure in the current pool to populate the next pool. Continue until the pool of currently predicted structures is empty. If, for a given currently predicted structure, the maximum allowed number of stems is reached (configurable parameter *maxstemnum*) or no stem that satisfies all thresholds can be added, the structure is moved from the pool to the set of predicted structures;
8. Repeat steps 0-7 for each defined parameter set;
9. Rank the predicted structures by their structure score, reactivity score, or the product of both scores (parameter rankby):

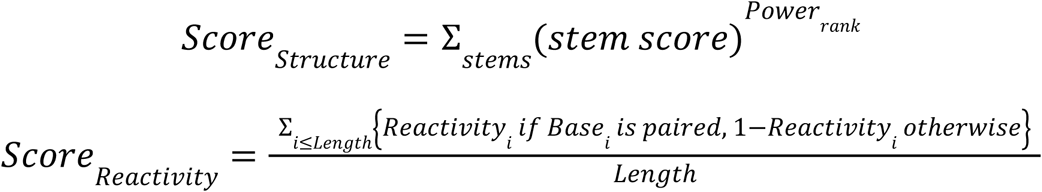

where *Power_rank_*is a scaling factor used to prioritize longer stems, and *Score_Reactivity_*is the MEA (Mean Absolute Error) value between the predicted structure and the input residue reactivities.

The *subopt* parameter is initialized to its starting value (*suboptmin*) and is increased by an increment value each time the size of the pool of currently predicted structures grows (in step 7) until the maximum defined value is reached (*suboptmax*). The increment value is determined by the number of increment steps (*suboptsteps*) as *(supoptmax - suboptmin) / suboptsteps*.

Any hard restraints defined by the user are considered in step 0, allowing only base pairs consistent with the restraints. In the case of multi-chain input, the distance factor is always set to 1 for stems confining chain breaks, and *(i, i + 1)*, *(i, i + 2)*, and *(i, i + 3)* base pairs are allowed for such stems. In the case of residue reactivities defined by the user, negative values of -10 and lower are treated as missing values, and the remaining values are clipped to the *0.0-1.0* range using *min(max(x,0),1)*. The slope (*M*) and intercept (*B*) parameters [58] are used to adjust the base pair probabilities with [RNA.fold_compound.sc_add_SHAPE_deigan()](https://www.tbi.univie.ac.at/RNA/ViennaRNA/doc/html/api_python.html) and to rescale the reactivity values such that *0.5* serves as a neutral (missing) value. The rescaled reactivities are then used to adjust the base pair weights, such that base pairs *(i, j)* with positive weights are multiplied by, and those with negative weights are divided by, the *reactivity factor:*

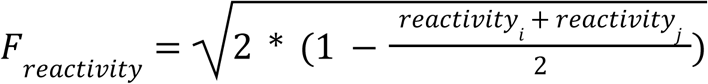

The theoretical time complexity of the greedy algorithm is *O(K * N^4^)*, where *N* is the sequence length and *K* is the number of predicted structures. Step 2 takes *O(N^2^)* time to annotate all stems, and step 3 takes *O(N)* time to score each of the *O(N^2^)* annotated stems, resulting in *O(N^3^)* time. Steps 2-6 are repeated *O(N)* times, corresponding to the number of stems in a predicted structure, resulting in an overall complexity of *O(N^4^)*. The algorithm runs relatively fast with *subopt = 1.0*, but slows down substantially at lower *subopt* values as the number of alternatives, *K,* grows exponentially.

The training procedure was performed on the <u>SRtrain150</u> dataset (the SRtrain entries shorter than 150nts in length, see the section on sequence datasets) through a series of grid search runs. The target metric was the F-score of the best structure among the top five predictions. After training, the following fixed values were selected: *Distance_ideal_* was set to 4 for hairpins and 2 for all other loops (distance factor formula); internal loops of the following sizes were defined as short near-symmetric loops deserving a bonus: *(0, 0), (0, 1), (1, 0), (1, 1), (0, 2), (2, 0), (2, 2), (1, 2), (2, 1), (3, 1), (1, 3), (2, 3), (3, 2), (3, 3), (3, 4), (4, 3), (4, 4), (4, 2), (2, 4)*; for symmetric loops *(x, x)* the bonus was doubled; for *(x - 1, x)* and *(x, x - 1)* loops the bonus was multiplied by 1.5; *Bonus_GNRA_* was set to 1.25; for the structure score the base pair weights were set to *GU = -1.0, AU = 1.5, GC = 4.0*, and the *Power_rank_* was set to 1.7. By default, no bifurcations were allowed once the pool of currently predicted structures reached 1000 structures (parameter *poollim = 1000*). For the Nussinov algorithm, the stem scores were used directly, whereas for the Edmonds and Hungarian algorithms, the stem scores were raised to the power of 1.7.

After training, the default configuration file (<u>def.conf</u>) included twelve parameter sets: **defG1** (algorithm G, *bpp = 0, bpweights: GC = 3.25, AU = 1.25, GU = -1.25, suboptmax = 0.9, suboptmin = 0.65, suboptsteps = 1, minlen = 2, minbpscore = 4.5, minfinscorefactor = 1.25, distcoef = 0.09, bracketweight = -2, orderpenalty = 1.0, loopbonus = 0.125, maxstemnum = 10^6^*), **defG2** (parameters different from defG1: bpweights: GC = 2, AU = 1, GU = 1, minbpscore = 3, minfinscorefactor = 0.99, distcoef = 0.1, orderpenalty = 1.35), **defN** (algorithm N, bpp = 0, bpweights: GC = 3.5, AU = 1, GU = -1, minlen = 2, minbpscore = 2.75), **bppN** algorithm N, *bpp = 0.5, bpweights: GC = 3.5, AU = 1, GU = -1, minlen = 2, minbpscore = 2.25*), **defE** (algorithm E, *bpp = 0, bpweights: GC = 3.75, AU = 1.75, GU = 0.5, minlen = 2, minbpscore = 4.5*), **defH** (algorithm H, *bpp = 0, bpweights: GC = 3.75, AU = 1.75, GU = 0.5, minlen = 2, minbpscore = 4.5*), **bppH1** (algorithm H, *bpp = 0.5, bpweights: GC = 4.0, AU = 0.5, GU = -1.0, minlen = 2, minbpscore = 2.25*), **bppH2** (algorithm H, *bpp = -1, bpweights: GC = 2.0, AU = 0.5, GU = -1.5, minlen = 2, minbpscore = 4.0*), **bppE1** (algorithm E, *bpp = 0.5, bpweights: GC = 4.0, AU = 0.5, GU = -1.0, minlen = 2, minbpscore = 2.25*), **bppE2** (algorithm E, *bpp = -1, bpweights: GC = 2.0, AU = 0.5, GU = -1.5, minlen = 2, minbpscore = 3.75*), **bppG1** (algorithm G, *bpp = -1.0, bpweights: GC = 2.0, AU = 0.5, GU = -0.5, minlen = 2, minbpscore = 3.25, minfinscorefactor = 1.25, orderpenalty = 0.5*), **bppG2** (algorithm G, *bpp = 0.5, bpweights: GC = 3, AU = 2, GU = 1, minlen = 2, minbpscore = 4.0, minfinscorefactor = 0.99, orderpenalty = 0.5*). Three of the configs (bppN, bppH1, bppH2) were assigned priority in the default structure ranking (parameter *priority*).

To imporve performance on longer RNA sequences, two faster configs were introduced, one for sequences longer than 500 nts (*<u>500.conf</u>*) and another for sequences longer than 1000 nts (*<u>1000.conf</u>*). Both differed from the default config only in the suboptimality-related thresholds (*suboptmin* and *suboptmax*), reducing the number of alternative structures considered. These configs were used for sequences of the corresponding lengths in all single-sequence benchmarks.

### SQUARNA algorithm for alignment-based prediction

For RNA secondary structure prediction from multiple sequence alignments, we developed a two-step procedure based on the principles of the single-sequence SQUARNA algorithm. Due to unclear stem boundaries in alignments caused by insertions and deletions, we adopted a base pair, rather than a stem, as the prediction unit. Step 1 consists of two iterations. In the first iteration, we calculate stem score matrices for each sequence in the alignment, sum them to obtain a total stem score matrix, and then greedily select a subset of compatible base pairs with the highest scores above a defined threshold. In the second iteration, the same procedure is performed while excluding base pairs incompatible with the previously selected subset. This iteration serves to refine the predicted structure by eliminating base pairs that belong to mutually incompatible stems with comparable scores. The base pairs selected after the second iteration constitute the predicted structure, referred to as SQUARNA step1. In step 2, we perform single-sequence SQUARNA predictions guided by the total stem score matrix obtained in step 1, iteration 1, and return the consensus of the individual predictions, referred to as SQUARNA step2. Step 2 helps identify base pairs that occur in the majority of sequences but do not reach the score threshold because of less-conserved neighboring base pairs. Finally, SQUARNA generates either the union (SQUARNA step3u) or the intersection (SQUARNA step3i) of the base pairs predicted in step 1 and step 2.

To compute the total stem score matrix in step 1, each sequence is first unaligned by removing all gaps. An individual stem score matrix is then computed using four parameters (*bpp, bpweights, minlen, minbpscore*). The resulting matrix is subsequently realigned to the alignment size by inserting all-zero rows and columns at gap positions in the aligned sequence. An individual structure for a given sequence in step 2 is defined as the consensus of the top-N (1 by default) ranked predicted structures based on a specified config. The overall consensus structure in step 2 is obtained by greedily selecting a consistent set of the most frequent (i.e., present in the largest number of sequences) base pairs from the individual structures that appear in at least a *freqlim* fraction of sequences. For both step 1 and step 2 predicted structures, a trimming procedure is applied that removes base pairs with pseudoknot orders exceeding a defined threshold (*levellim*). In step 3, the union of step 1 and step 2 base pairs (in the *step3u* setting) is computed by adding consistent step 2 base pairs to the base pairs obtained at step 1.

The parameter training procedure was performed on the RNAStralignExt dataset (see the section on alignment datasets). The target metric was the F-score of the SQUARNA *step3u* setting. After training, the following parameter values were selected: *levellim = 3* for alignment lengths under 500 nts and 2 otherwise; *freqlim* was set to 0.35; step1 parameters (from <u>ali.conf</u>): *bpp = 0, bpweights GC = 3.25, AU = 2.0, GU = -1.0, minlen = 2, minbpscore = 4.5*; the config for the single-sequence SQUARNA predictions at step 2 (*<u>ali.conf</u>*) consisted of a single parameter set, **ali**, with the following values: algorithm G, *bpp = 0, bpweights GC = 3.25, AU = 2.0, GU = -1.0; suboptmax = 1.0; suboptmin = 1.0; suboptsteps = 1; minlen = 2; minbpscore = 4.5; minfinscorefactor = 1.0; distcoef = 0.09; bracketweight = -2; orderpenalty = 0.75; loopbonus = 0.125; maxstemnum = 10^6^*. At step 2, the base pair score matrix in the individual predictions was multiplied by the normalized total stem score matrix obtained at iteration 1 of step 1. For this purpose, the total stem score matrix was normalized to a maximum value of 5.0 (by dividing all values by the maximum and then scaling to 5.0). The normalized stem score matrix was also unaligned separately for each sequence to match the correct dimensions.

The theoretical time complexity of SQUARNA step 1 is *O(D * N^2^)*, where *D* is the alignment depth (number of sequences) and *N* is the alignment length, as the computation of each of the *D* individual stem score matrices takes *O(N^2^)* time to annotate all potential stems. In the benchmarks, SQUARNA step1 was the second-fastest algorithm after RNAalifold. The theoretical time complexity of SQUARNA step 2 is *O(D * K * N^4^)*, corresponding to the complexity of the single-sequence SQUARNA multiplied by the number of sequences in the alignment.

### SQUARNA tool implementation

SQUARNA was implemented in Python 3 and is available as a Python library (<u>pip install</u> <u>SQUARNA</u>), as a command-line tool (https://github.com/febos/SQUARNA), and as a web server (https://larnal.imol.institute/). Single-sequence predictions were parallelized either over the pool of currently predicted structures (default) or over the input sequences (*byseq* option). Alignment-based predictions were parallelized over aligned sequences by default. Single-sequence predictions with alignment input were implemented using an unalign-realign procedure to account for gaps. In SQUARNA, we used a custom implementation of the Nussinov algorithm, a SciPy implementation of the Hungarian algorithm (scipy.optimize.linear_sum_assignment), and a NetworkX implementation of the Edmonds algorithm (networkx.algorithms.matching.max_weight_matching). The tool supports various input file formats, including FASTA, Stockholm, and Clustal, as well as a custom FASTA-like format, enabling the incorporation of restraints, reactivities, and reference structure data.

Three automated searches for structural restraints were implemented, available for single input sequences: Rfam family templates (*rfam* option), G-quadruplex patterns (*g4* option), and RNA-binding protein (RBP) binding motifs (*rbp* option). The Rfam family search uses the cmscan utility from the Infernal package [11], with the options “-E 1e-4” (E-value threshold), “--rfam” (Rfam-like score thresholds), and “--toponly” (ignore reverse strand). The cmscan-predicted structures are subsequently mapped onto the input sequence in decreasing order of significance, while ignoring inconsistent base pairs. For G-quadruplexes, based on recent benchmarks [28, 29, 67], we employ a regular expression search for two patterns, G_2-5_N_1-2_G_2-5_N_1-2_G_2-5_N_1-2_G_2-5_ and G_3-5_N_1-12_G_3-5_N_1-12_G_3-5_N_1-12_G_3-5_, followed by filtering using a G4Hunter score threshold of 1.2 [67]. For RBP-binding motifs, we performed a literature search and identified six well-defined sequence motifs: AUUGCAC (all unpaired, U1A [68]), GAAACAC (GC base pair flanking a loop of unpaired residues, Fab [69]), GGAGA (all unpaired, LIN28 [70]), UGCAUG (all unpaired, RBFOX1/2 [71]), UGUAHAUW (H = A/C/U, W = A/U, all unpaired, PUM [72]), ACUAAC (all unpaired, SF1/QKI [73]). The web server was implemented using React (frontend) and Django (backend) frameworks and is designed specifically for the analysis of individual RNA sequences, with all automated searches for restraints enabled by default. RFviewJS (https://github.com/dincarnato/RFviewJS) and RNAcanvas [74] were used for 2D visualizations.

### RNA sequence datasets

The <u>SRtrain</u> dataset was prepared as follows. RNA 3D structures from the PDB [36] were selected according to a representative set of RNA structures [75] (version 3.278 with a 3.0 Å cutoff). RNA secondary structures were annotated using DSSR [76] (version 2.0). Subsequently, a custom Python script was used to extract RNA strands from PDB entries, subject to three conditions: (i) the strand is continuous, without chain breaks or missing residues; (ii) the strand does not contain modified residues; and (iii) the strand does not form WC or wobble base pairs with any other strand, but forms at least one such base pair within the strand. The resulting set of strands underwent manual secondary structure redundancy reduction, yielding a non-redundant subset of 274 RNA strands with corresponding secondary structures. The SRtrain150 subset included 266 strands shorter than 150 nts.

The <u>SRtest</u> dataset was collected using the same procedure from another version of the representative set of RNA structures [75] (version 3.322 without a resolution cutoff). Sequences with at least 80% identity to any sequence in the SRtrain dataset were excluded. The resulting set of strands underwent manual secondary structure redundancy reduction, yielding a non-redundant subset of 247 RNA strands with corresponding secondary structures.

The <u>S01</u> dataset was prepared as follows. A set of 24 sequences, along with SHAPE data, was downloaded from [26] (https://webshare.oasis.unc.edu/weeksgroup/data-files/ShapeKnots_DATA.zip). RNA secondary structure data were converted from .ct format to dot-bracket notation. The original SHAPE data contained values ranging from 0.0 to 1.5, with some outliers and missing values encoded as -999. Two sequences required adjustments: (i) for the entry “5’ domain of 16S rRNA, H. volcanii” one excess value at the 3’-end was removed; (ii) for the entry “SARS corona virus pseudoknot” three missing -999 values were added at the 3’-end. The supplementary information from [26] (https://www.pnas.org/doi/suppl/10.1073/pnas.1219988110/suppl_file/sd01.pdf) was used as a reference for these fixes.

### RNA multiple sequence alignment datasets

The <u>RNAStralignExt</u>, <u>RfamPDB</u>, and <u>Rfam14.9</u> datasets were prepared by obtaining seed alignments in <u>Stockholm</u> format from Rfam [10] (version 14.9, 4,108 families). The Stockholm files were then converted into one-line <u>FASTA</u>-like and <u>Clustal</u> formats using custom Python scripts. The 134 families of the RfamPDB dataset were identified via a search for “Families with 3D structure” on the Rfam website (https://rfam.org/search?q=entry_type:%22Family%22%20AND%20has_3d_structure:%22Yes%22). All benchmarks for Rfam14.9 and RfamPDB were performed with the exclusion of the eleven RNAStralignExt families.

The <u>SubAli</u> dataset was prepared as follows. For each of the eleven Rfam families in the RNAStralignExt dataset, ten alignments were randomly sampled at each of the eleven depths (where permitted by the seed alignment depth): 2, 5, 10, 20, 30, 50, 75, 100, 150, 200, and 300 sequences. The sampled sequences were not subjected to any realignment procedures. The <u>SeqSim</u> dataset, used to analyze the correlation between prediction quality and mean sequence similarity of the alignment, was prepared as follows. For each of the eleven Rfam families in the RNAStralignExt dataset, 100 alignments were randomly sampled at each of three depths - 5, 10, and 20 sequences (where permitted by the seed alignment depth). For each alignment of a given (family, depth) pair, the mean pairwise sequence similarity was computed and rounded to three decimal places. Subsequently, only one alignment for each (family, depth, similarity) triple was retained, resulting in approximately 50 distinct alignments for each (family, depth) pair. The Hamming distance (https://en.wikipedia.org/wiki/Hamming_distance) between aligned sequences was used as the measure of pairwise sequence similarity. The sampled sequences were not subjected to any realignment procedures.

### Benchmarks

To assess prediction quality, the number of true positives (*TP*) was defined as the number of correctly predicted base pairs, the number of false positives (*FP*) as the number of incorrectly predicted base pairs, and the number of false negatives (*FN*) as the number of missed base pairs. The number of true negatives (*TN*) was not defined. Precision was defined as *Precision = TP / (TP + FP)*, recall as *Recall = TP / (TP + FN)*, and the F-score as the harmonic mean of precision and recall, *F-score = 2 * Precision * Recall / (Precision + Recall) = 2 * TP / (2 * TP + FP + FN)*. The F-score served as the main evaluation metric. In all benchmarks, inconsistently predicted base pairs (non-GC/AU/GU pairs and *(i, i + k)* pairs with *k < 4)* were removed prior to evaluation. The benchmarks were conducted on an AMD Ryzen 9 5950X machine with 32 CPU cores and 128 GB RAM.

For single-sequence benchmarks, we used IterativeHFold [45], HotKnots [46], IPknot [15], CONTRAfold [47], Knotty [48], EternaFold [49], RNAfold [24], ShapeKnots [50], Fold [50], CentroidFold [51], ProbKnot [50], CParty [52], HFold [53], RNAsubopt [24], RibonanzaNet-SS [54], MXfold2 [19], SPOT-RNA [20], eFold [22], and UFold [55]. For alignment-input benchmarks, we used CentroidAlifold [18], RNAalifold [24], R-scape [57], and IPknot [15]. SPOT-RNA2 [21] was not included in the benchmarks, as it takes a single-sequence input and constructs an alignment of homologous sequences on the fly; consequently, it’s unclear to which tool category it should belong.

**IterativeHFold** was obtained as part of the Shapify package (version 1.0) from https://github.com/ltrinity/Shapify and executed as *“HFold_iterative --s ‘SEQUENCE’”* (single prediction) and *“HFold_iterative --s ‘SEQUENCE’ --n 5”* (five predictions). **HotKnots** (version 2.0) was obtained from https://www.cs.ubc.ca/labs/algorithms/Software/HotKnots/ and executed as *“HotKnots -s SEQUENCE”*. We were unable to obtain HotKnots predictions for the two longest RNA sequences in the SRtest dataset. **IPknot**, obtained from https://github.com/satoken/ipknot (version 1.0.0), was executed as *“ipknot inputfile > outputfile”*. For the SeqSim dataset, IPknot exceeded ten days of runtime on the alignment RF00177_5_729.sto, and for the Rfam14.9 dataset, it exceeded ten days on the RF02746 family alignment; consequently, IPknot benchmarks for these two datasets were omitted. **CONTRAfold**, obtained from http://contra.stanford.edu/contrafold/download.html (version 2.02), was executed as *“contrafold predict inputfile > outputfile”*. **Knotty**, obtained from https://github.com/TheCOBRALab/Knotty (version 1.0.0), was executed as *“knotty SEQUENCE > outputfile”*. We were unable to obtain Knotty predictions for the three longest RNA sequences in the SRtest dataset. **EternaFold**, obtained from https://github.com/eternagame/EternaFold (version 1.3.1), was executed as *“contrafold predict inputfile --params ./EternaFold-1.3.1/parameters/EternaFoldParams.v1 > outputfile”*. The **RNAfold**, **RNAsubopt**, and **RNAalifold** tools were obtained as part of the ViennaRNA package (version 2.7.0) and executed as *“RNAfold --noPS [--shape=shapefile] inputfile > outputfile”*, *“RNAsubopt --sorted [--shape=shapefile] inputfile > outputfile”*, and *“RNAalifold --noPS inputfile [--shape=shapefile] > outputfile”*. Due to RAM limitations, the *“--deltaEnergy=0.1”* setting was used to run RNAsubopt for sequences longer than 2000 nts. **ShapeKnots**, **ProbKnot**, and **Fold** were obtained as part of the RNAstructure package (version 6.5) and executed as *“DATAPATH=../data_tables ./ShapeKnots-smp inputfile outputfile [-sh shapefile]”*, *“DATAPATH=../data_tables ./ProbKnot inputfile outputfile --sequence”*, and *“DATAPATH=../data_tables ./Fold inputfile outputfile”*. **CentroidFold** and **CentroidAlifold**, obtained from https://github.com/satoken/centroid-rna-package (version v0.0.16), were executed as *“centroid_fold inputfile > outputfile”* and *“centroid_alifold inputfile > outputfile”*. Additionally, we used *“centroid_fold -g 6 inputfile > outputfile”* (CentroidFold(g=6)), as we observed improved performance compared to the default setting. **CParty**, obtained from https://github.com/TheCOBRALab/CParty (version 1.0.0), was executed as *“CParty SEQUENCE > outputfile”* (single prediction) and *“CParty -n 5 SEQUENCE > outputfile”* (five predictions). **HFold**, obtained from https://github.com/TheCOBRALab/HFold (version 1.0.2), was executed as “*HFold SEQUENCE > outputfile*” (single prediction) and “*HFold -n 5 SEQUENCE > outputfile*” (five predictions). **RibonanzaNet-SS** was obtained from https://www.kaggle.com/code/shujun717/ribonanzanet-2d-structure-inference (version 20) and executed as a Jupyter Notebook. **MXfold2**, obtained from https://github.com/mxfold/mxfold2 (version 0.1.2), was executed as *“mxfold2 predict inputfile > outputfile”*. **SPOT-RNA**, obtained from https://github.com/jaswindersingh2/SPOT-RNA (as of April 16, 2023), was executed as *“python SPOT-RNA.py --inputs inputfile --outputs outputfolder --cpu 32”*. **eFold** was obtained via *“pip install efold”* (version 0.1.2) and executed as *“efold SEQUENCE -o outputfile”*. **UFold**, obtained from https://github.com/uci-cbcl/UFold (version 1.2), was executed as *“python ufold_predict.py”*. We were unable to obtain UFold predictions for six sequences in the SRtest dataset (lengths: 8, 9, 10, 11, 17, and 20 residues). **R-scape**, obtained from https://github.com/EddyRivasLab/R-scape/tree/master/versions/rscape (version 2.5.6), was executed as *“R-scape --fold --covmin 4 --nofigures --rna inputfile > outputfile”*. The *“Rscape nested”* structure was defined as the *“SS_cons”* line of the output, and the *“Rscape total”* structure was defined as the *“SS_cons”* line with the addition of consistent base pairs from the *“SS_cons_N”* lines of the output. To identify significantly covarying base pairs, we run R-scape with the command *“R-scape --nofigures --rna inputfile > outputfile”* and used the default E-value threshold of 0.05. Version 3.22 of SQUARNA was used in all benchmarks.

## DATA AVAILABILITY

The benchmarking data are available at https://github.com/febos/SQUARNA-data (DOI: https://doi.org/10.5281/zenodo.20533263).

## CODE AVAILABILITY

The SQUARNA code is available at https://github.com/febos/SQUARNA (DOI: https://doi.org/10.5281/zenodo.8292325).

## ACKNOWLEDGMENTS

The authors thank Danny Incarnato, Sean Eddy, Maciej Cieśla, and all members of the Bujnicki group at IIMCB for their valuable feedback and discussions. E.F.B. thanks his former supervisor Mikhail Roytberg for the visionary idea of using a stem as the prediction unit, and Pavel Pevzner and Phillip Compeau for their online course “Bioinformatics Algorithms: An Active Learning Approach”, which had a major impact on this work.

## FUNDING

E.F.B. and M.D.S. were supported by the National Science Centre, Poland [NCN grant 2024/53/B/NZ2/02718 to E.F.B.]. J.M.B. and G.I.N were supported by the National Science Centre, Poland [NCN grant 2017/26/A/NZ1/01083 to J.M.B.]. E.F.B. was additionally supported by the European Molecular Biology Organization [EMBO fellowship ALTF 525-2022 to E.F.B].

## AUTHOR CONTRIBUTIONS

E.F.B. designed the computational method. E.F.B., M.D.S., D.R.B., and G.I.N. developed the computational tool. All authors performed the analyses and edited the manuscript. E.F.B. and J.M.B. supervised the work.

## COMPETING INTERESTS

The authors declare no competing interests.

## SUPPLEMENTARY MATERIALS

**Supplementary Figure S1.**
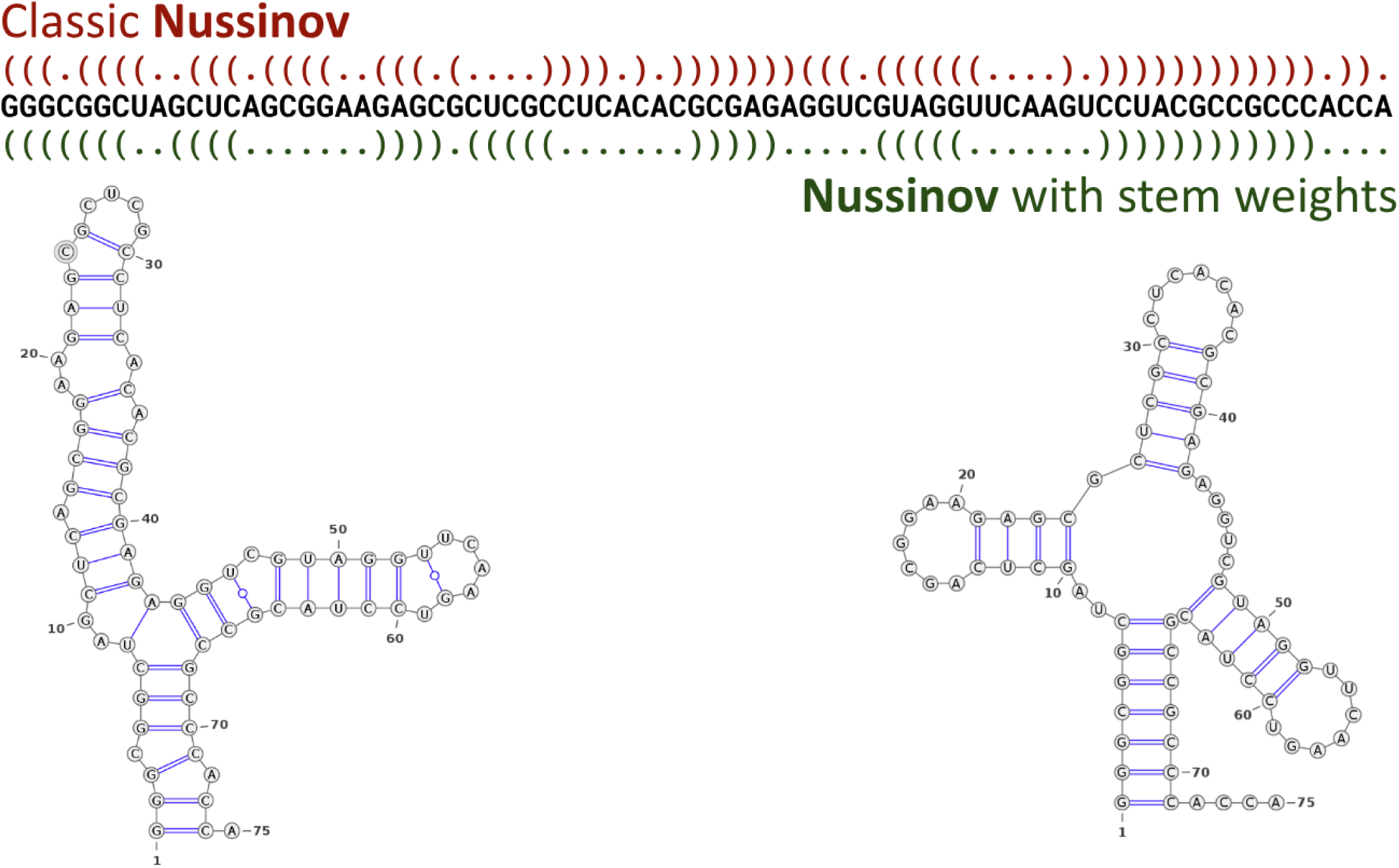
Performance of the Nussinov algorithm on a tRNA sequence. The structure on the left and in red was predicted by the classical Nussinov algorithm, with G-C, A-U, and G-U base pairs each assigned a score of 1. The structure on the right and in green was predicted by a Nussinov variant using a stem weight matrix, with base pair scores of 3 for G-C, 2 for A-U, and 1 for G-U. Structure visualizations were generated using the VARNA applet (https://varna.lisn.upsaclay.fr/).

**Supplementary Figure S2.**
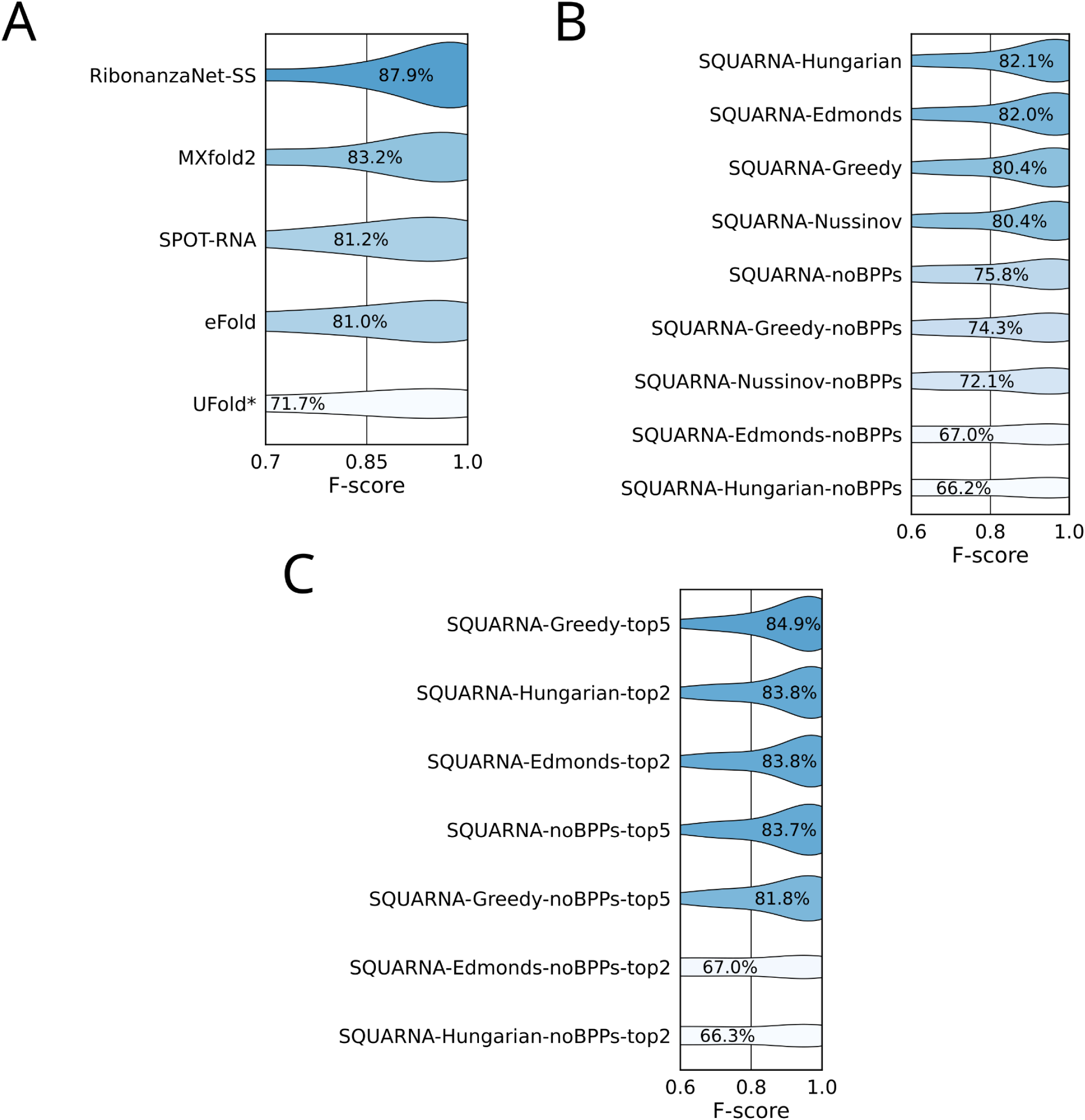
Single-sequence SQUARNA performance: additional plots. (**A**) Violin plot of F-score values achieved by deep learning models on the SRtest dataset of 247 sequences, considering only the top-ranked predictions. (**B**) Violin plot of F-score values achieved by several SQUARNA variants on the SRtest dataset of 247 sequences, considering only the top-ranked predictions. (**C**) Violin plot of F-score values achieved by several SQUARNA variants on the SRtest dataset of 247 sequences, considering the best of the top five predictions. Asterisks denote tools that could not process all sequences in the dataset and were assigned F-scores of 0.0 for those sequences, see the Methods section for details; “top2” denotes SQUARNA variants that report at most two predicted structures; “Hungarian”, “Edmonds”, “Nussinov”, and “Greedy” denote SQUARNA variants that use only the corresponding algorithm; “noBPPs” denotes SQUARNA variants that do not utilize base pair probabilities derived with ViennaRNA [**ViennaRNA**].

**Supplementary Figure S3.**
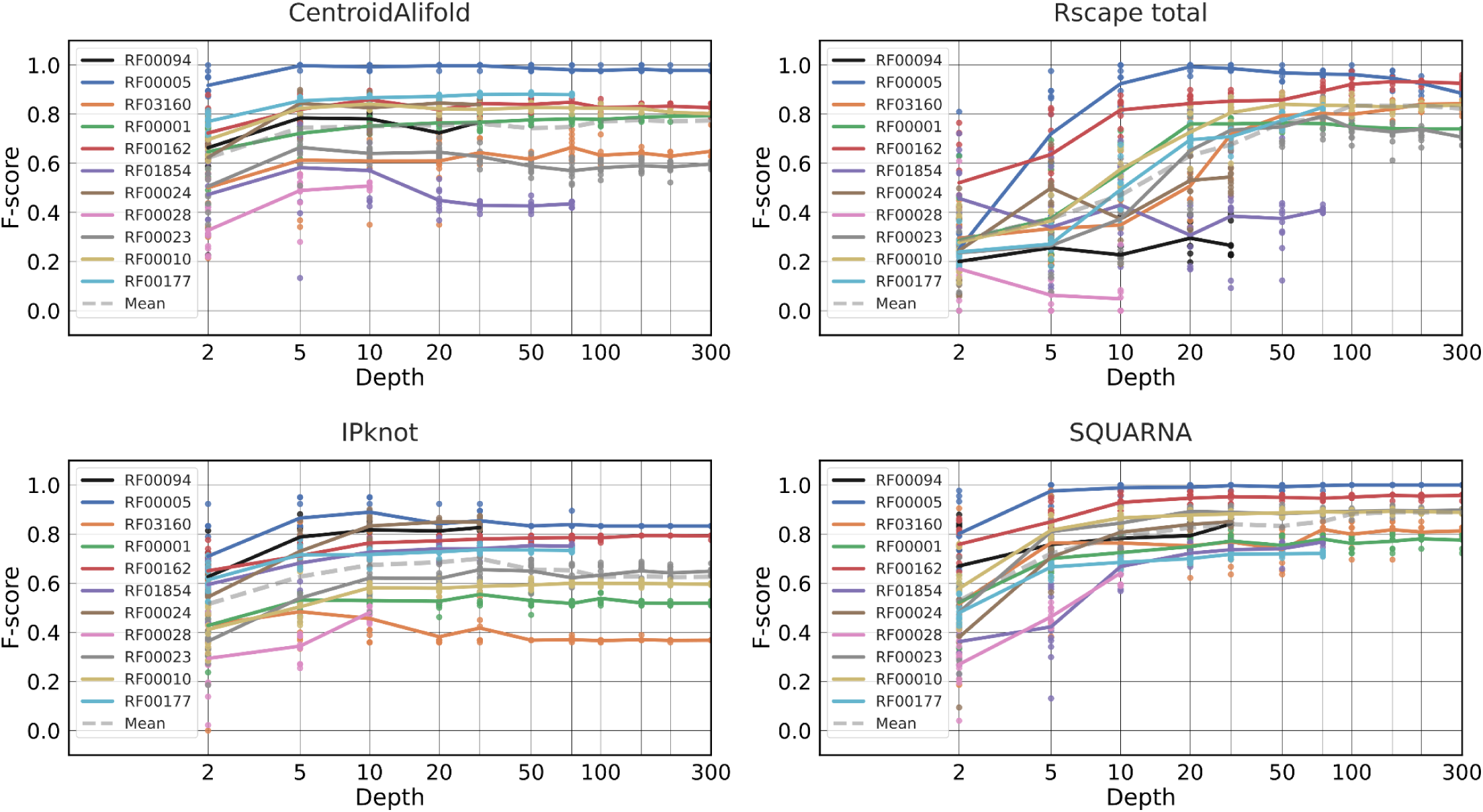
Alignment-based SQUARNA performance: additional plots. Prediction quality of CentroidAlifold, Rscape, IPknot, and SQUARNA on the SubAli dataset. Each dot represents the F-score of a single prediction, and solid lines depict F-scores averaged over the ten alignments of each depth for a given Rfam family. The dashed gray line indicates the mean F-score across all eleven Rfam families.

**Supplementary Figure S4.**
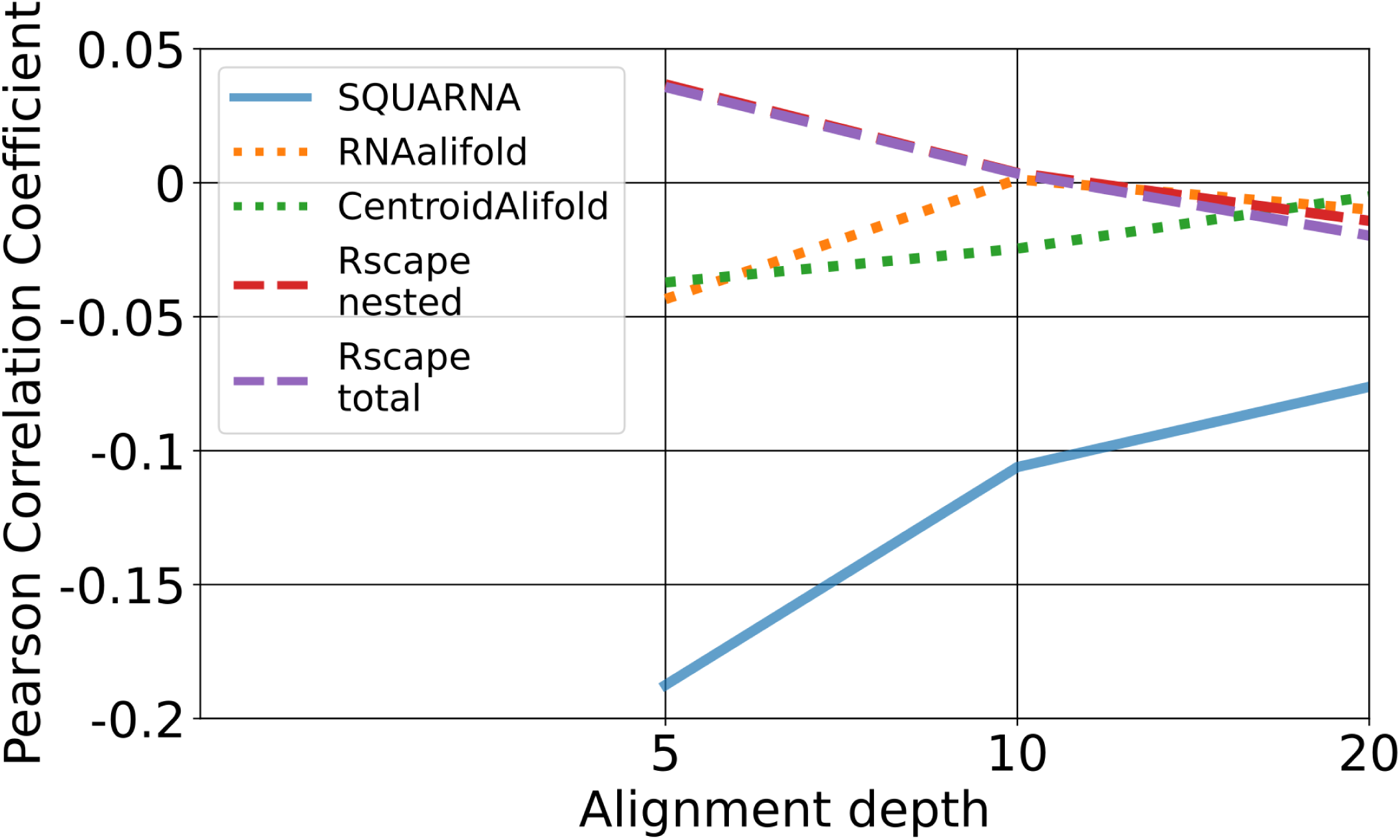
Influence of sequence similarity on prediction quality. Pearson correlation coefficients between the prediction quality of alignment-based tools, measured as F-scores, and the mean pairwise sequence similarity of the input alignment, plotted as a function of alignment depth. The benchmark was performed on the SeqSim dataset, see the Methods section for details. SQUARNA exhibits a modest negative correlation, whereas the other tools show near-zero correlation values.

**Supplementary Figure S5.**
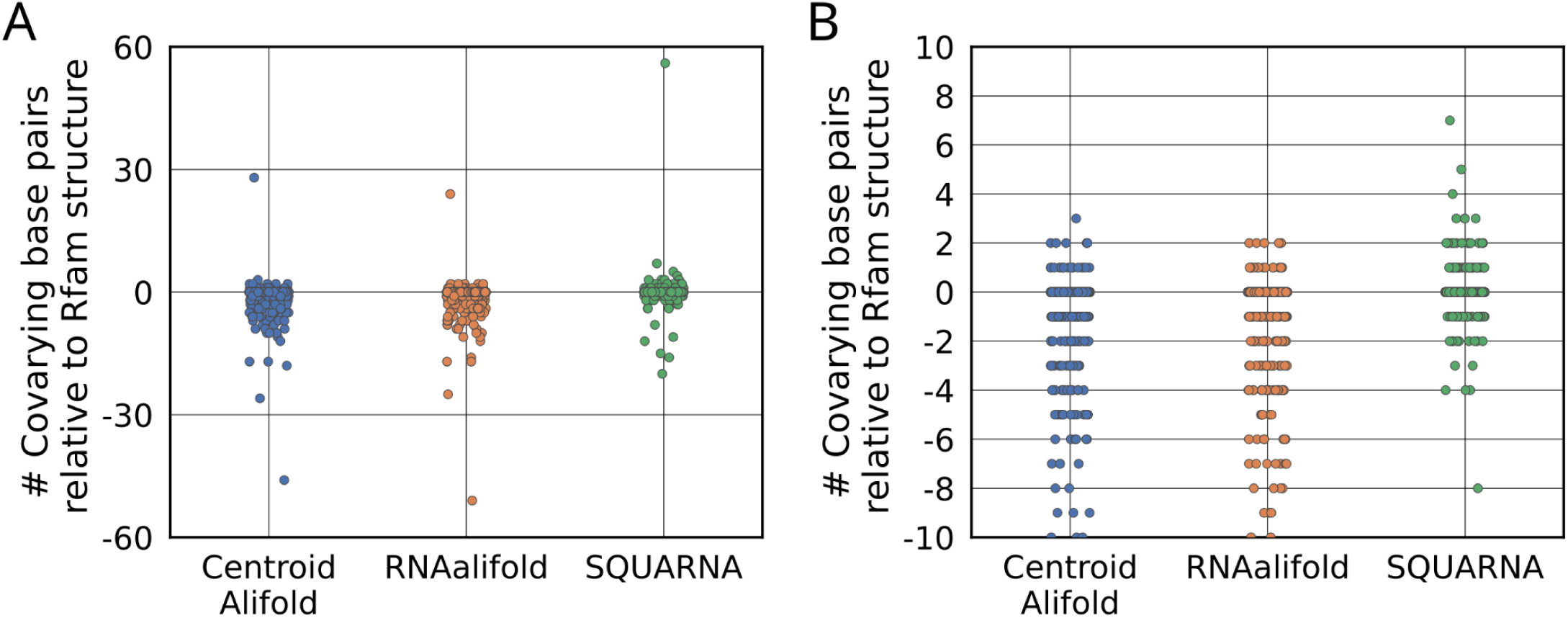
Benchmarking results in terms of covarying base pairs. Benchmarking results showing the number of significantly covarying base pairs, as identified by R-scape, among the base pairs predicted by CentroidAlifold, RNAalifold, and SQUARNA. The Y-axis shows the difference between each tool’s prediction and the Rfam-annotated structure for 1,011 families from the Rfam 14.9 dataset in which at least one significantly covarying base pair was identified by R-scape. Each dot represents a single Rfam family. (**A**) Outliers included. (**B**) Outliers excluded.

**Supplementary Figure S6.**
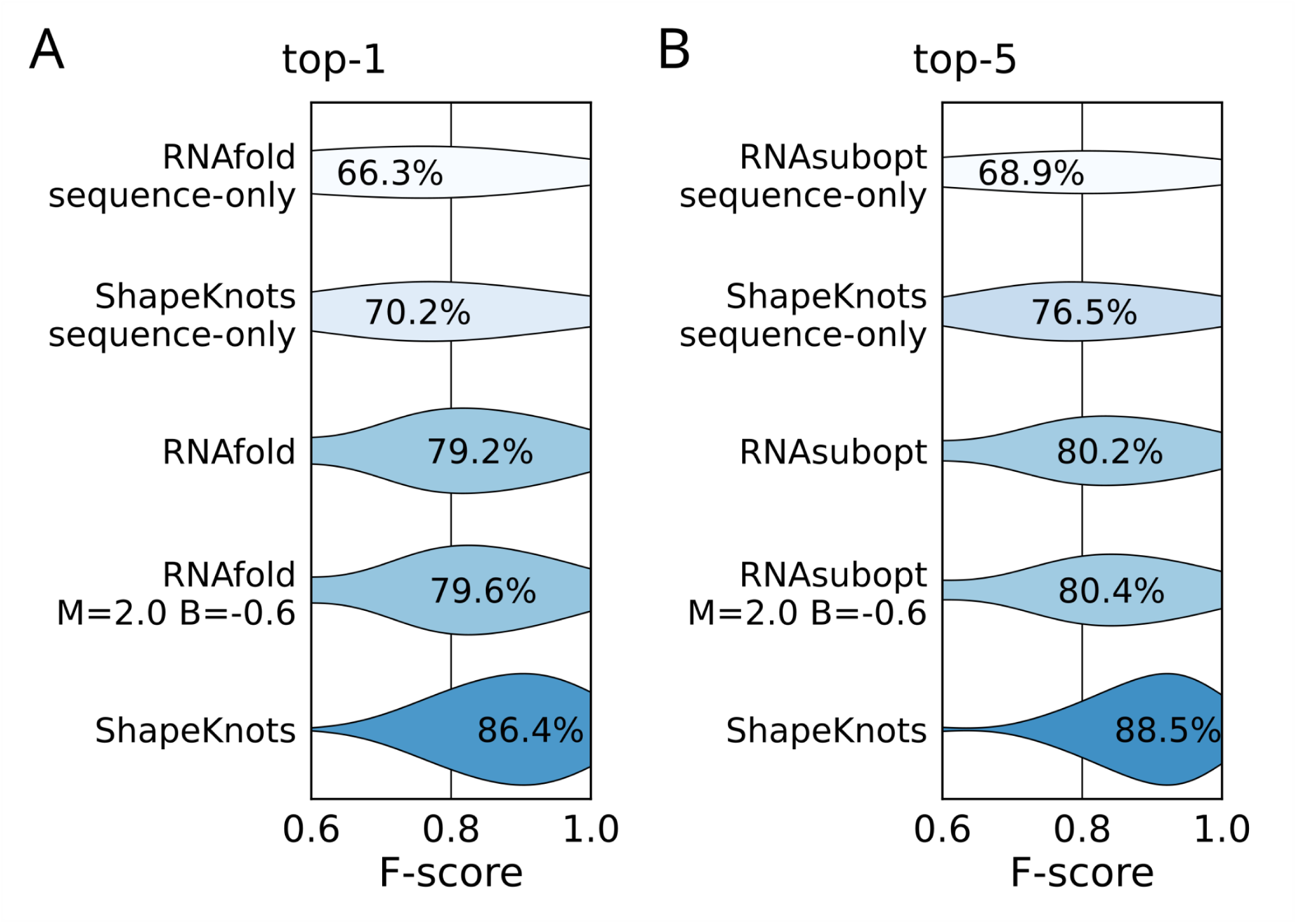
Benchmarks with input residue reactivities. Distributions of F-scores measured for (**A**) the top-ranked prediction and (**B**) the best of the top five predictions generated by existing tools for 24 RNA sequences from the S01 dataset. The “sequence-only” variants do not use residue reactivities and consider only sequence input. The “M=2.0 B=-0.6” variant shows the performance of RNAfold and RNAsubopt using optimized M and B values obtained for the S01 dataset through grid search.

**Supplementary Table S1.** Statistics of stem formation detected in the SRtrain and SRtest datasets derived from RNA-containing PDB entries, 521 sequences in total. Each counted stem represents a potential non-extendable stem with at least one base pair formed in the annotated structure. Each such stem was assigned a base pair mask with 1 indicating a present base pair and 0 indicating an absent base pair. Thus, 1111 denotes a 4bp stem that matches its non-extendable version, 11110 denotes a 5bp stem lacking one flanking base pair, and 1011011 denotes a partially formed 7bp stem belonging to the “Other” category.

|  | 3bp stems | 4bp stems | 5bp stems | 6bp stems | 7bp stems | >7bp stems | total |
| --- | --- | --- | --- | --- | --- | --- | --- |
| **Non-extendable** | 100%<br>(352) | 73.8%<br>(313) | 62.9%<br>(282) | 58.4%<br>(181) | 52.4%<br>(118) | 46.5%<br>(177) | 66.5%<br>(1423) |
| **Missing one flanking bp** | 0 | 26.2%<br>(111) | 21.9%<br>(98) | 26.4%<br>(82) | 25.8%<br>(58) | 21.8%<br>(83) | 20.2%<br>(432) |
| **Other** | 0 | 0 | 15.2%<br>(68) | 15.2%<br>(47) | 21.8%<br>(49) | 31.7%<br>(121) | 13.3%<br>(285) |
| **Total** | 352 | 424 | 448 | 310 | 225 | 381 | 2140 |

**Supplementary Table S2.** Contents of the RNAStralignExt dataset. Additional families with pseudoknotted structures are highlighted by underlining.

| **RNAStralignExt dataset** |  |  |  |  |
| --- | --- | --- | --- | --- |
| **N** | **Family** | **Rfam ID** | **Length** | **Depth** |
| 1 | 5S rRNA | RF00001 | 230 | 712 |
| 2 | tRNA | RF00005 | 118 | 954 |
| 3 | RNaseP | RF00010 | 996 | 458 |
| 4 | tmRNA | RF00023 | 954 | 477 |
| 5 | telomerase RNA | RF00024 | 777 | 37 |
| 6 | group I intron | RF00028 | 892 | 12 |
| 7 | <u>HDV ribozyme</u> | <u>RF00094</u> | 115 | 33 |
| 8 | <u>SAM riboswitch</u> | <u>RF00162</u> | 239 | 457 |
| 9 | 16S rRNA | RF00177 | 1980 | 99 |
| 10 | SRP RNA | RF01854 | 315 | 92 |
| 11 | <u>Twister ribozyme</u> | <u>RF03160</u> | 189 | 1613 |

**Supplementary Table S3.** Distribution of Rfam 14.9 alignments with predicted secondary structures by tool (as indicated in the “#GF SS” field of the Stockholm-format seed alignment file).

| **Tool \ Depth** | **2 seq** | **3 seq** | **4 seq** | **5 seq** | **>5 seq** | **total** |
| --- | --- | --- | --- | --- | --- | --- |
| **R-scape** | 1 | 3 | 2 | 1 | 2 | 9 |
| **LocARNA** | 1 | 1 | 1 | 0 | 15 | 18 |
| **mfold** | 49 | 25 | 11 | 7 | 25 | 117 |
| **PFOLD** | 2 | 2 | 6 | 25 | 206 | 241 |
| **RNAalifold** | 771 | 287 | 188 | 122 | 490 | 1858 |
| **Total** | 1006 | 520 | 355 | 283 | 1944 | 4108 |

## Notes

https://github.com/febos/SQUARNA

https://github.com/febos/SQUARNA-data

https://larnal.imol.institute/

### Competing Interest Statement

The authors have declared no competing interest.

### Summary of Updates

Revised title and figures (improved styles, no data changes)

